# Cascades of neuronal plasticity in the macaque visual cortex

**DOI:** 10.1101/2022.09.17.508292

**Authors:** Kenji W. Koyano, Elena M. Esch, Julie J. Hong, Elena N. Waidmann, David A. Leopold

**Author notes:** These authors contributed equally to this work (Elena M. Esch, Julie J. Hong, Elena N. Waidmann). Medical Scientist Training Program, University of Colorado Anschutz Medical Campus, Aurora, CO 80045. Laboratory of Neurogenetics of Language, The Rockefeller University, 1230 York Avenue, New York, NY, 10065.

## Abstract

Primates readily learn new visual objects, with neurons in the inferior temporal cortex exhibiting diminished visual responses to familiar stimuli. The mechanisms of visual plasticity expressed within a neural population are largely unknown. Here we used chronic microwire electrodes in a face-selective cortical area in the macaque to longitudinally track cohorts of neurons across several weeks of exposure to novel faces. Neurons showed gradual adaptation in their late-phase visual responses, with cell-specific time constants ranging from two to twenty days. These time constants were governed by the number of testing days rather than by the cumulative number of stimulus exposures. This gradual buildup of altered visual responses may serve as an internal and graded mark of stimulus familiarity, a central component of visual recognition.

## Main Text

The neocortex of the mammalian brain processes complex sensory information (*1*) and stores memory for subsequent recognition across specialized circuits in the brain, including the cerebral cortex (*2, 3*). Primates have an advanced capacity for the visual comprehension of objects and social stimuli, which is reflected in specialized visual regions of the inferior temporal (IT) cortex (*4, 5*). Neurons in IT exhibit experience-dependent response plasticity following repeated exposure to the same visual stimuli (*6, 7*), for example showing diminished responses to familiar visual objects compared to novel ones, particularly in the late-phase of the response (*8-13*). Despite these advances, methodological restrictions have limited our understanding of important mechanistic elements of visual response plasticity, such as the time course of individual neurons and population dynamics more broadly.

Here we examined the unfolding of neural plasticity in the inferior temporal cortex of the macaque by tracking the responses of isolated single neurons during up to five weeks of daily exposure to visual stimuli. During this period, we randomly interleaved presentations of 120 visual stimuli that were initially unfamiliar to the animals and 60 stimuli that were already highly familiar. Longitudinal tracking with a flexible microwire electrode array allowed us to observe how neurons gradually acquired the memory for individual stimuli across recording days (Fig. 1A, B). Prior to any visual exposure to the stimuli, the electrodes were surgically implanted into the fMRI-defined anterior medial (AM) face patch (Fig. 1A,C,D), the most anteroventral node in the macaque IT face processing network, which sits adjacent to the perirhinal cortex and is thought to play an important role in processing facial identity (*14*). The microwires permitted isolation and tracking of the same neurons during the experimental sessions over weeks (Fig. 1E, Fig. S1) (*15-17*).

**Figure 1.**
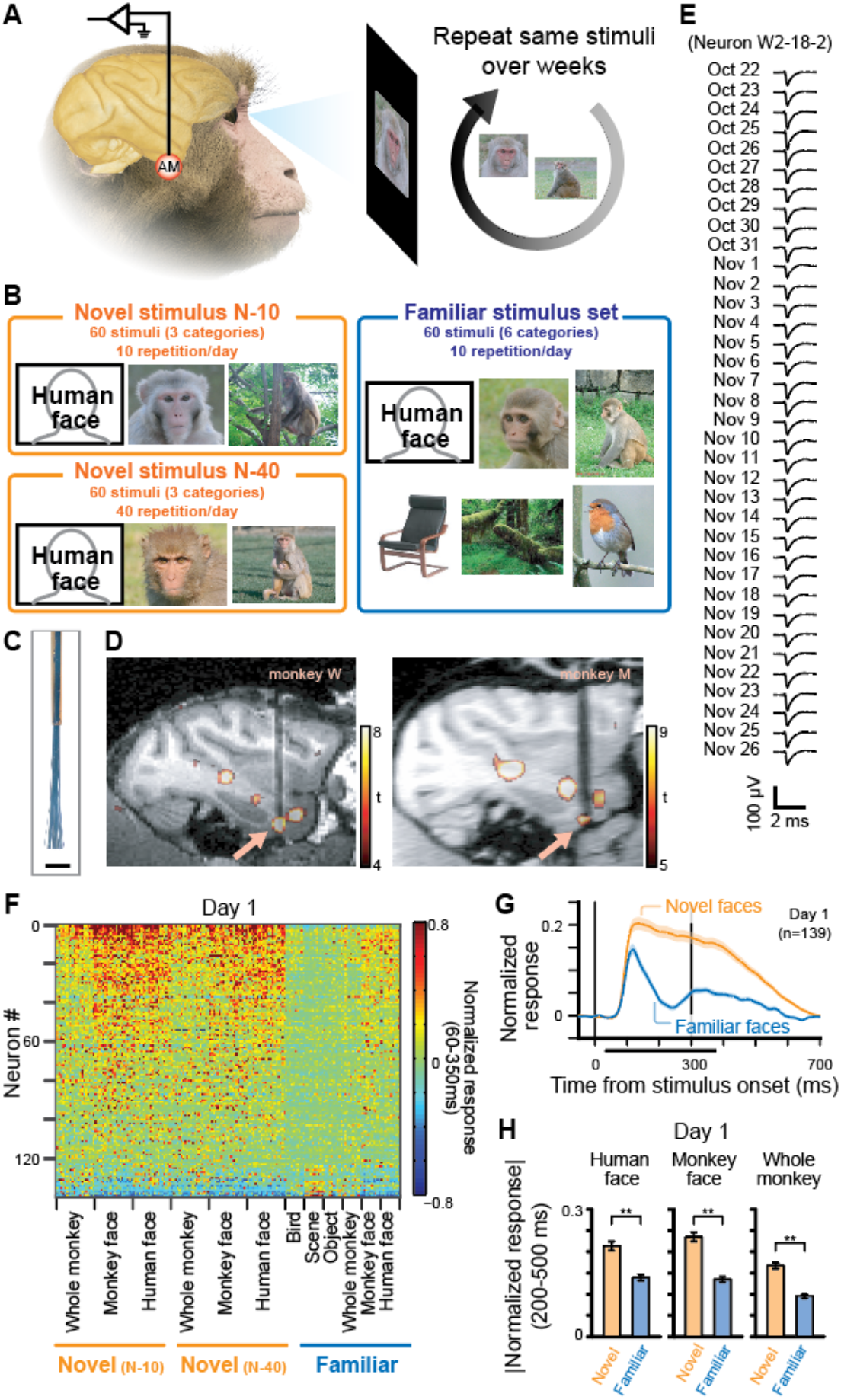
Experimental design and population response to the stimuli. **A**, Overview of the experiment. Neurons were recorded from AM face patch while stimuli were repeatedly shown to the monkeys over the course of 2-5 weeks. **B**, Three categories (novel N-10, novel N-40 and familiar) of visual stimuli. Human face images are substituted to line drawing, according to bioRxiv policy. **C**, Electrode tip of the microwire brush array. Scale bar, 1 mm. **D**, Recording location of each monkey, shown in sagittal section of MRI. Arrows, AM face patch. **E**, Example waveform of a neuron recorded over 36 days. **F**, Response of recorded neurons to each stimulus on the first day of the recording. Neurons were sorted according to the response to face stimuli compared to other stimuli. **G**, Time course of responses to novel and familiar face stimuli on the first day. Shaded area indicates standard error from mean (S.E.). **H**, Average response to each category of stimulus on the first day during the late sustained response. Ordinate, absolute value of normalized responses. Error bars, S.E. **p<10^−14^, paired t-test.

We carried out longitudinal recordings from the AM face patch from two animals through four extended experimental sessions, each ranging from 11 to 37 days of continuous recording. Throughout these sessions, we isolated a total of 4874 neural waveforms over 90 days of recordings (54.8 ± 5.4 isolated neurons per day). We restricted analysis to 139 neurons that were longitudinally recorded for at least 7 days (Fig. S2) and responsive to at least one stimulus in each of novel and familiar categories (p<0.05, t-test versus baseline response with Bonferroni correction). Of 139 neurons, 117 (84.2%) were face-selective (face-selective index>0.33, see Methods and Fig. S3).

Consistent with previous reports, population responses on the first experimental day were larger to novel stimuli than familiar stimuli (Fig. 1F), primarily during the late sustained response (Fig. 1G, see Fig. S4) (*9-13*). Similar patterns of late-phase response suppression were observed for familiar images of human faces, monkey faces, and whole monkeys (Fig. 1H).

Fig. 2A shows the responses of an example neuron to four stimuli recorded daily over a period of five weeks, starting on October 22. Two of the stimuli were initially novel and then presented ten times per day. The other two were highly familiar (presented 685 times through 48 days; see Methods) and also presented ten times per day. The novel stimuli at first elicited sustained spike trains, lasting well beyond the removal of the stimulus. However, these sustained responses decreased very gradually, over the course of several days, until they were no longer detectable after 2.5 weeks, around November 10 (Fig. 2A, *left two columns*). The early visual response between 100 and 200 ms after the stimulus onset was comparatively unchanged during this period and still evident on November 26, after five weeks of exposure to the novel stimuli.

**Figure 2.**
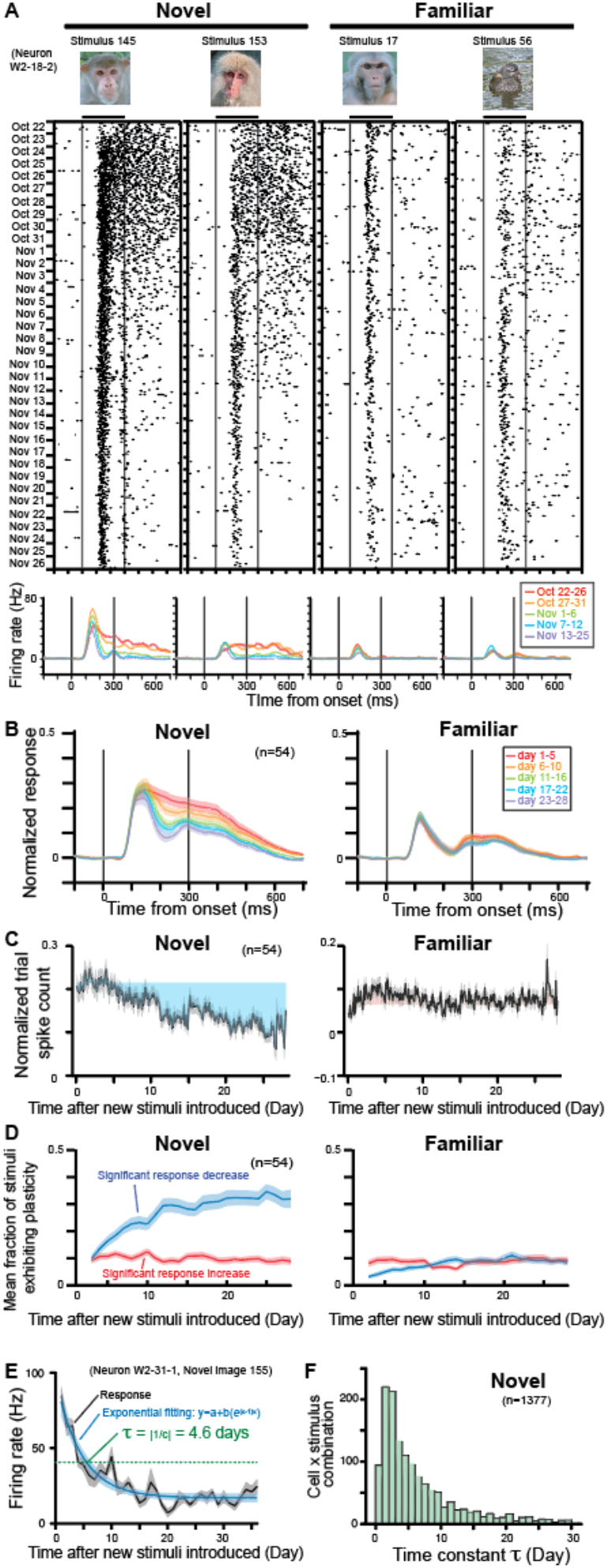
Decrease of late, sustained responses over multiple days of visual exposure to initially novel stimuli. **A**, Response of an example neuron over five weeks of recording, showing adapting responses to novel (left) but not familiar (right) stimuli. **B**, Population-averaged spike density functions for novel (N-10) and familiar stimuli. **C**, Change of response during late sustained period of 200-500 ms from stimulus onset over four weeks. All presentation trials are plotted in order, with days indicated at bottom. Gray shaded area, S.E. from mean. Colored shaded area, difference from mean response of first two days. **D**, Mean fraction of stimuli showing significant population response changes from first two days (t-test, p<0.05). **E**, An example of exponential fitting and computation of τ for response changes for a given neuron x stimulus combination across the extended recording session. **F**, Distribution of time constants τ for all neuron x stimulus combinations in the novel (N-10) stimulus set.

Responses of the same neuron to familiar stimuli were much weaker relative to the novel stimuli beginning on the first experimental day, a finding that was representative across the population (Fig. 1F-H), and relatively constant through the five weeks (Fig. 2A, *right two columns*).

To examine these response changes across the population of neurons, we analyzed recording sessions that were conducted over a period of greater than four weeks in each monkey.

Across a population of 54 neurons recorded at least 20 of the first 28 days (mean recording period = 30.8 ± 5.4 days), a gradual decline in late-phase responses was clearly evident (Fig. 2B, C, see also Fig. S5 for other example neurons). Analysis revealed that the sustained responses to novel stimuli decreased significantly (Fig. 2B, C, *left*. p<10^−6^, t^46^=5.85, paired t-test between first and last two days), while the corresponding responses to familiar stimuli did not (Fig. 2B, C, *right*, p=0.25, paired t-test, t^46^=1.16). The fraction of novel stimuli whose population responses were affected by this form of experience-dependent plasticity grew to reach 31.9 ± 3.3% after four weeks (Fig. 2D, *left*, mean ± S.E., see also Fig. S6). In contrast, the fraction of familiar stimuli affected was minimal (Fig. 2D, *right*). While most changes were expressed during the sustained period, early transient responses did show a small response increase for a subset of novel stimuli (11.4 ± 1.6% at four weeks later, mean ± S.E., Fig. S7). The observed response change could not be attributed to non-specific factors, such as a general adaptation of the cell or a loss of spike isolation over time, since a new set of novel stimuli introduced in the middle of the experimental session initially elicited strong sustained responses, which subsequently declined over time (Fig. S8).

To quantify the time constant of the response decay following the introduction of novel stimuli, we fit the responses of each neuron to each stimulus with an exponential function (Fig. 2E), matching the observed response decay and eventual attainment of a stable value. We restricted our analysis to responses from 1377 neuron × stimulus combinations that could be fit well with an exponential through the course of the session of 11-36 days (see Methods for detailed criteria). As only a subset of stimuli showed this behavior for a given neuron, this criterion limited analysis to 16.5% of all possible neuron × stimulus combinations. We defined the time constant τ as the time point when the neuron’s response decreased to 1/e, or 36.8 %, of the response on the first day (green dotted line in Fig. 2E). Across the 1377 combinations, the distribution of τ had a long tail, ranging from 2 days to nearly 30 days. The median τ value was 4.23 days (Fig. 2F), after which responses are estimated to decrease by 95% after 12.7 days. This range of time constants for stimulus familiarity is broadly consistent with estimations from previous studies tracking multi-unit responses (*18*) and responses of different neurons (*19*) across days. Quantifying the time constant of individual IT neurons enabled us to further evaluate important factors for the time course and population dynamics as described in following sections.

The response to novel stimuli decreased with repeated daily exposure to the same stimuli. Would these changes proceed faster if more repetitions were presented each day? We tested this question by conducting an additional longitudinal experiment in both animals comparing the time constant between two novel stimulus sets, one shown 10 times per day (N-10 stimulus set) and the other shown 40 times per day (N-40 stimulus set). Fig. 3A shows the responses of an example cell for two stimuli, including one from N-10 and one from the N-40 stimulus sets. Both sets of stimuli show gradually decreasing sustained responses over weeks. To weigh the relative contribution of total days versus total number of presentations, we computed both time-and trial-based exponential models for the N-10 and N-40 stimuli (Fig. 3B).

**Figure 3.**
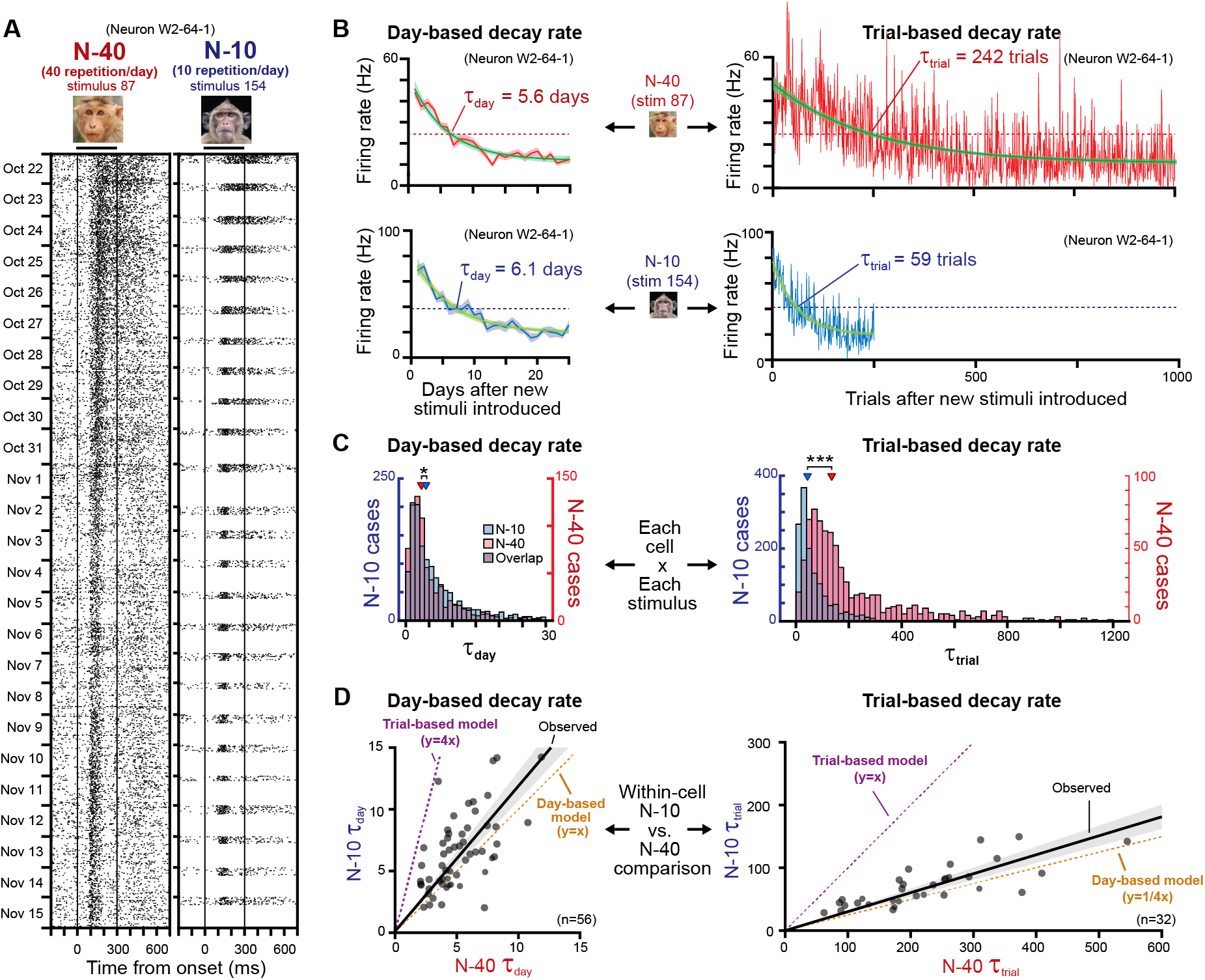
Primary factor of day on the rate of visual response plasticity. **A**, Response change of an example neuron for one of N-40 and N-10 stimuli. **B**, Exponential fitting used to calculate day-based (*left*) and trial-based (*right*) decay rates. Upper two plots, example for the N-40 stimulus. Bottom two plots, example for the N-10 stimulus. The good correspondence between day-based decay rates and divergent trial-based decay rates for the N-40 and N-10 stimuli indicate that day is the critical factor. **C**, Population distribution of estimated decay rates for each cell x stimulus combination. *Left*, day-based decay rate (τ_day_). *Right*, trial-based decay rate (τ_trial_). *p<10^−5^, ***p<10^−122^, Mann-Whitney U test. **D**, Within-cell comparison between N-10 and N-40 stimuli. Each dot represents a cell. The black regression line is calculated from observed τ with a linear regression model y=ax.

This analysis revealed that the number of days determined the rate of responses change to a much higher degree than did the number of presentations. In this example, this can be seen in the similar day-based time constant (τ_day_, Fig. 3B, left) but strongly divergent trial-based time constant (τ_trial_, Fig. 3B, right).

Across the population, the number of days, rather than the accrued number of stimulus presentations, were the critical determinant of response adaptation rate (Fig. 3C). For the N-40 and N-10 stimulus sets, the overall distributions of τ_day_ for all neuron x stimulus combinations was highly similar, though there was a small difference (median 3.36 vs. 4.32 days, p<10^−5^, Mann-Whitney U test). In contrast, the median distributions for τ_trial_ differed approximately by a factor of four (median 135.2 vs. 43.6 trials, p<10^−122^, Mann-Whitney U test). Comparing the mean time constants of individual neurons of both animals for the N-10 and N-40 stimuli similarly indicated that day was the critical variable (Fig. 3D). Neurons had very similar τ_day_ values for the two conditions and were thus distributed near the unity line (Figure 3D, *left*, black regression line, R^2^ = 0.55, slope of 1.18); however, τ_trial_ values that differed by approximately nearly a factor of four (Figure 3D, *right*, black regression line, R^2^ = 0.69, slope of 0.30). These results indicate that the number of elapsed days during periods of visual exposure critically determines the rate of response plasticity among single neurons.

Most of the analysis thus far considered the temporal dynamics of plasticity regarding all possible neuron x stimulus combinations. Given the broad distribution of well-modeled time constants, we next asked whether the rate of plasticity is set by individual neurons or by individual stimuli. For example, if each neuron has a unique and fixed time constant, then a given neuron should show the same rate of plastic changes for multiple different stimuli. On the other hand, if the rate of plasticity is determined principally by details of the stimulus, then the entire neural population should adjust its activity in concert. In that case, a given neuron could exhibit vastly different plasticity time constants for different stimuli.

We thus analyzed the plasticity time constants as a function of both neurons and stimuli (Fig. 4). We found that neural identity had a much stronger role in determining the rate of plasticity than did stimulus identity. For individual cells, the plasticity rate was comparable for different stimuli; however, for a stimulus, the plasticity rate widely across different neurons (Fig. 4A, see also Fig. S9 for divergent plasticity rate for individual stimulus). Across the population of the 1377 neuron × stimulus combinations, cell identity had a very strong and significant effect on the time constant (Fig. 4B, p<10^−33^, Kruskal-Wallis test). The main effect of cell identity was robust and reproduced in each experimental session (p<0.02 -p<10^−8^, Fig. S9) and for the N-40 stimuli (p<10^−29^).

**Figure 4.**
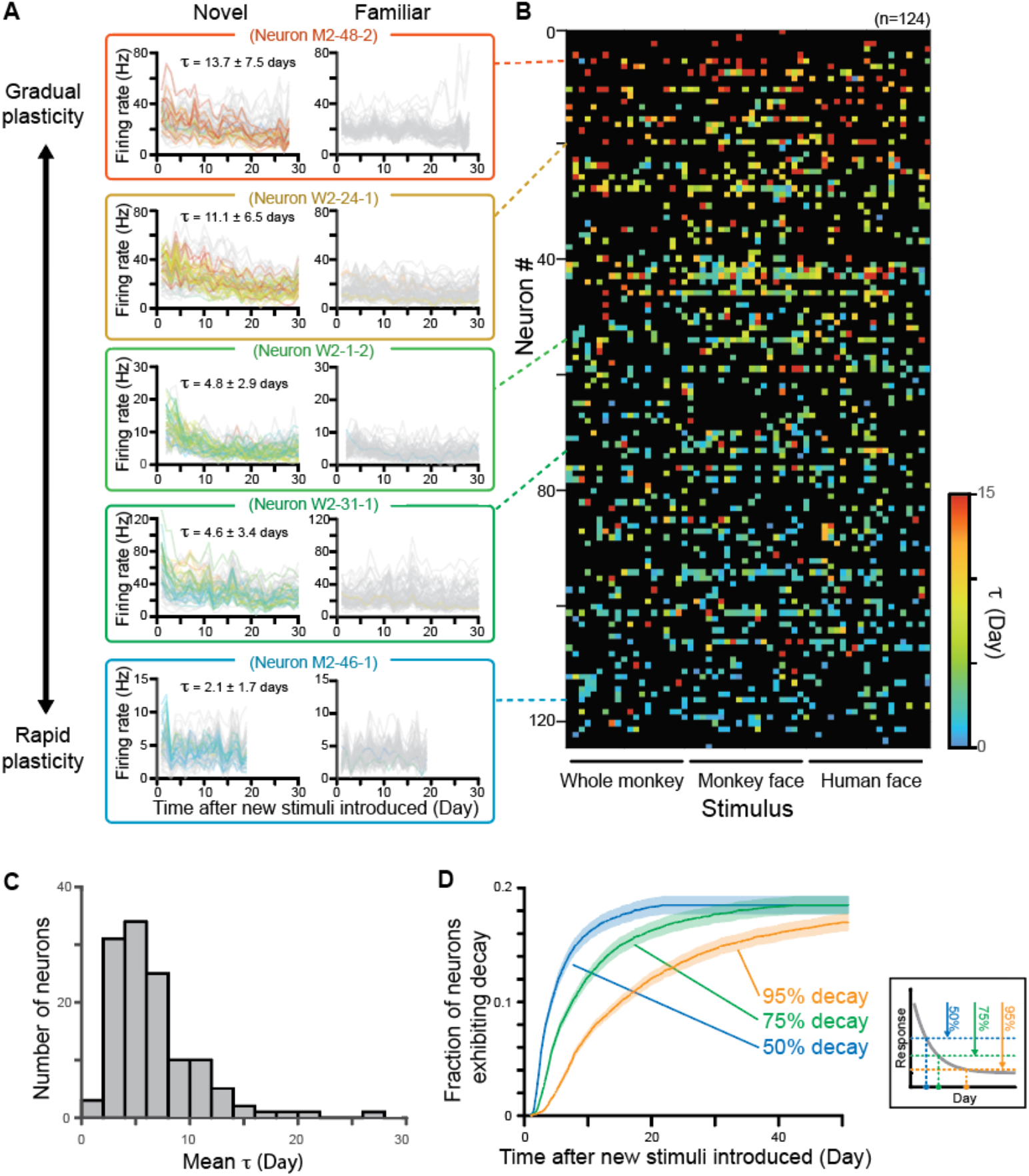
Cascades of neuronal plasticity during the repeated presentation of stimuli across weeks. **A**, Example neurons highlighting the wide range of plasticity time constants across the population. The colors indicate the τ_day_ for each stimulus depicted in the color scale in **B**. Gray traces correspond to stimuli that did not elicit significant plasticity. Neurons were sorted with average time constant, from gradual plasticity neurons (top) to rapid plasticity neurons (bottom). **B**, Time constant for all cell x stimulus combinations across the population, showing dependence of the length of time constant on neuron’s identity. Black squares correspond to stimuli that do not exhibited change of responses or do not meet the criteria of fitting quality. Neurons had similar time constants for multiple stimuli, whereas individual stimuli did not show shared time constants across neurons. **C**, Distribution of mean τ of each neuron. **D**, Ratio of neurons exceeding 50, 75 and 95% decay threshold over days, which is estimated from τ of each neuron x stimulus combination. The fraction of neurons is calculated for each stimulus and then averaged. Inset, definition of time when a neuron exceeds decay thresholds.

These results indicate that the population of neurons in the AM face patch have a broad range of unique cell-specific time constants governing their plasticity. These time constants ranged from two days to more than twenty days (Fig. 4C). Following the introduction of new stimuli, the number of neurons showing modified responses to the stimuli thus increases over time. Fig. 4D shows that, depending on the criterion for response modification, the proportion of participating neurons grows gradually over days and weeks. In the case of the AM face patch, nearly 20% of the recorded neurons, on average, eventually exhibited plasticity to the face and body stimuli we showed.

The observation that individual IT neurons have unique time constants for visual plasticity suggests a division of labor that may bear on the graded nature of familiarity in visual recognition (*20, 21*). The brain’s internal signal for stimulus familiarity is thought to be continuous and to increase with increasing exposure. Our findings suggest that visual face-selective neurons may contribute to this internal signal through a progression of commitment. According to this hypothesis, the most rapidly adapting neurons we observed would be specialized to identify and remember new experiences, perhaps opening the door to the recruitment of additional neurons following further exposure. With increasing experience playing out across days, neurons with longer plasticity time constants joined the subpopulation of affected neurons. Thus, over time, and through this observed cascade of neural plasticity, the population stores an increasingly intense mark of familiarity that could serve visual recognition.

The psychological variable of familiarity is most frequently associated with the perirhinal cortex (*8, 19, 22-25*), and has recently been associated with an adjacent region in the temporal pole (*26*). Nonetheless, the evidence for involvement of IT cortex in this process is also strong (*8-13*). For example, IT neurons are more selective (*10, 13*), respond more reliably (*12*) and show sharper dynamic responses (*11*) for familiar stimuli, suggesting that experience has the capacity to shape high-level sensory processing in favor of known stimuli.

The face patch AM from which we recorded is directly adjacent to the perirhinal cortex and is most commonly associated with the encoding of individual face identity (*14, 27*). In AM and several other lateral and ventral face patches, Landi and Freiwald found diminished fMRI activation for visually familiar versus non-familiar faces (*25*). By contrast, face patches far away from this region, such as those in the superior temporal sulcus (STS) showed weak if any effects of familiarity. Perhaps for this reason, an earlier longitudinal recording from the anterior fundus (AF) face patch showed little evidence of response changes over several months (*15, 16*). Face patches in these areas may be principally concerned with dynamic aspects of facial behavior and may therefore offer less contribution to visual recognition (*28-30*). Whether attending to changing aspects of faces, such as motion, gaze and expression, contributes to long-term plasticity in such areas is a question for the future.

Primates are highly social animal that form large groups and interact competitively with other groups. Accurately learning and retaining knowledge about individuals, including their facial structure and behavior, is a critical aspect of daily survival. Our findings that AM neurons in the adult brain exhibit progressive plasticity for faces may tie directly to primates’ lifelong capacity to learn new individuals, or to improve their capacity to read and interpret the faces of their conspecifics. It is notable that the plasticity observed in the present study was during the late response period, similar to the late-period responses involved in analyzing the details of facial identity (*31*) or in discounting average facial structure in theories of norm-based encoding (*32, 33*). It is possible that plasticity of late-period responses in the AM face patch additionally serves the continual updating of an internal model of facial statistics to aid in identifying individuals or interpreting facial behavior. Like the familiarity signal, this process may benefit from the gradual commitment of neurons over visual experience and may thus share overlapping neural mechanisms with the gradual familiarization of novel stimuli studied here. For example, during the learning process, rapidly adapting neurons can quickly modulate their tuning for new identities, amid a larger population of more stable neurons that maintains codes for reliable recognition. The mixture of stable and flexible information may be a universal property of adaptable systems in the brain, as similar principles appear to be present in other structures, such as the basal ganglia (*34*).

## Materials and Methods

### Subjects

Two rhesus monkeys (macaca mulatta, both male, monkey W and M weighing 8.5 and 9.3 kg, respectively) were used in this study. All animals were surgically implanted with an MRI-compatible head post, and with a chronic microwire electrode bundle in an AM face patch (Fig. 1D) which was functionally localized using a standard fMRI block design using movie clips (*35, 36*) and/or a naturalistic movie watching paradigm (*37*). The apparatus and surgical implantation protocol have been described in detail previously (*16*). All surgeries were performed under aseptic conditions and general anesthesia under isoflurane, and animals were given postsurgical analgesics and prophylactic antibiotics. During participation in the recording experiment, the animals were on water control and received their daily fluid intake during their testing (see below). Each subject’s weight and hydration level were monitored closely and maintained throughout the experimental testing phases. All the experimental procedures and animal welfare were in full compliance with the Guidelines for the Care and Use of Laboratory Animals by U.S. National Institutes of Health and approved by the Animal Care and Use Committee of the U.S. National Institutes of Mental Health / National Institute of Health.

### Behavioral task and visual stimuli

The animals were not required to perform a behavioral task that required cognitive efforts for recognition of identities. With the passive viewing paradigm, neurons were examined for sensory processing of visual face stimuli. The monkeys sat in a primate chair in front of an LCD/OLED monitor with their head position stabilized by means of an implanted head post. They were required to maintain their gaze on a fixation point of 0.2° × 0.2° at the center of the monitor through a trial. In each trial, visual stimuli of 15° diagonal length (7.3-12.9° × 7.6-13.1°) were presented for 300 ms in pseudo-random order followed by a 400 ms inter-stimulus interval. The monkeys were rewarded with fruit juice for successfully maintaining fixation within a window of 1.5° −2°, while their eye position was monitored using an infrared video-tracking system (EyeLink II; SR Research). Stimulus presentation, eye position monitoring, and reward delivery were controlled by MonkeyLogic software (*38*) or NIMH MonkeyLogic Software (*39*). The monitor (either a ViewSonic 18” LCD monitor or LG 55” OLED monitor) was placed 90 cm in front of the monkey. Timing of stimulus presentation was recorded by a photodiode sensor that received signal from a small white square displayed on a corner of the screen at the same time of stimulus presentation.

The main stimulus set comprised three groups of stimuli, each of which included 60 images (Fig. 1B). One stimulus set was highly familiar to the animals and the other two stimulus sets were novel to the animals at the beginning of each series of experimental session. The familiar stimulus set includes images from 6 categories: human face, monkey face, whole monkey, object, scene, and bird (10 images from each category). Each of the two novel stimulus sets includes images from three categories: human face, monkey face and whole monkey (20 images from each category). The human face images were drawn from the FEI face database (*40*), and monkey faces and bodies were provided courtesy of Dr. Olga Dal Monte. Images from all other stimulus categories were assembled from web searches or iPhone applications (*16*). The images of the familiar stimulus set had been used to confirm neurons’ consistent responses across days (also see Electrophysiology section below) in other studies and repeatedly presented to the animals before starting the first data collection of this study. For monkey M, the familiar stimuli had been presented 68,948 times in total (mean: 1,149 times per each stimulus) through 65 experimental days before the first data collection of this study. For monkey W, the familiar stimuli had been presented 41,102 times in total (mean: 685 times per each stimulus) through 48 experimental days before the first data collection of this study. We continued to use the same familiar images during data collection in this study. The images of the novel stimulus sets had never been shown to the animals until starting data collection for this study, and we prepared new stimuli when starting new series of experimental session. The familiar stimulus set and one of the novel stimulus sets (N-10) were shown to the animals 10 times per day, while the other novel stimulus set (N-40) was shown 40 time per day. In addition to the three major stimulus sets described above, another “intermittent” novel stimulus set was also shown to the animals 10 times on the first day and on the eleventh or twelves day. In one series of experimental session (session W-2), this intermittent stimulus set was also shown to the animal on days 19, 26, 33, 36 and 37. The detailed stimulus presentation schedule is shown in Supplementary Figure 2. There are total of four series of experimental sessions, each of which continued for 11, 11, 28 and 37 days, respectively. In two longer series of experimental sessions, (sessions M-2 and W-2), additional new novel stimulus sets (N-10_2_, N-10_3_ and N-10_4_) were introduced at the middle of the experimental sessions, on days 13, 19 and 26. These additional novel stimulus sets were shown 10 times a day. To keep the total number of image presentation per day the same, 15 images were randomly dropped from N40 stimulus set each time when the new stimulus set was introduced. In one of the recording sessions (W-1), familiar and intermittent stimulus sets were presented one day before and one day after the series of the experimental session only, since the animal could not complete larger number of trials during that time.

### Electrophysiology

Extracellular neuronal signals were recorded with 64 chronically implanted NiCr wires (Microprobes, Fig. 1C) that permitted tracking of individual neurons over multiple recording sessions (*15-17*). The recorded neuronal signals were amplified and digitized at 24.4 kHz in a radio frequency-shielded room by PZ5 NeuroDigitizer (Tucker-Davis Technologies), and then stored to an RS4 Data Streamer controlled by an RZ2 BioAmp Processor (Tucker-Davis Technologies). A gold wire inserted into a skull screw was used for ground. Broadband signals (2.5-8 kHz) were collected from which individual spikes were extracted offline using the WaveClus software (*41*) after filtering between 300 and 5000 Hz. Event codes, eye positions and a photodiode signal were also stored to a hard disk using OpenEX software (Tucker Davis Technologies).

The method for longitudinal identification of neurons across days was described in detail previously (*15, 16*). The spikes recorded from the same channel on different days routinely had closely matching waveforms and interspike interval histograms and were provisionally inferred to arise from the same neurons across days. This initial classification based purely on waveform features and spike statistics was further tested against the pattern of stimulus selectivity and temporal structure of the neurons’ firing evoked by visual stimulation with the familiar stimulus set (Supplementary Figure 1). We used the distinctive visual response pattern to the familiar stimuli generated by isolated spikes as a neural “fingerprint” to further disambiguate the identity of single units over time (*32, 42*). The stability of stimulus selectivity was assessed by calculating a correlation coefficient for two firing rate vectors for the 60 familiar stimuli between two consecutive days (Supplementary Figure 1 D, G). In addition, the consistency was confirmed by the stability of firing rate strength during the baseline period (−100 to +50 ms relative to stimulus onset, Supplementary Figure 1E) and mean response to the familiar stimuli (60 to 350 ms from stimulus onset, Supplementary Figure 1G). For subsequent analysis we used 139 neurons which were isolated at least for 7 days (Supplementary Figure 2). Some neurons were isolated thought a series of recording sessions, while some other neurons disappear, emerge and reappear during the recording session.

### QUANTIFICATION AND STATISTICAL ANALYSIS

Stored neuronal response data were analyzed offline with MATLAB software (Mathworks, MA). All the data in the text was expressed as mean ± S.D. unless otherwise stated. Error bars in figures are standard error unless otherwise stated. All the t-test statistics were two-tailed. When a neuron was not isolated on a given day and data is missing, we perform statistics without the data (Fig. 2B, seven neurons were not isolated on the first or last two days and excluded from t-test; Supplementary Fig. 1E, three neurons were not isolated on the first or last two days in session W-1 and excluded from t-test). Firing rate responses of neurons to each stimulus were calculated for the following periods: baseline period, 150 ms before the stimulus onset to 50 ms after stimulus onset; response period, 60 to 350 ms after the stimulus onset; late sustained period, 200 to 500 ms after the stimulus onset; early transient response period, 50 to 150 ms after the stimulus onset. Significant responses to each stimulus were evaluated by t-test with Bonferroni correction for firing rate responses between baseline and response period. All the 139 neurons showed significant response to at least one stimulus of each stimulus sets of familiar and novel stimuli and were considered as responsive to the stimuli. Population-averaged tuning was calculated by averaging normalized firing rate response, which was calculated from the maximum and baseline responses of each neuron. Maximum response was defined as the response to the stimulus that elicited the largest response during the response period, either excitation or suppression, as compared to the baseline response. The response for each stimulus was normalized by subtracting the baseline response and then dividing it by the absolute difference between the baseline and maximum response of the neuron, resulting in normalized firing rate value that ranges from −1 to 1 (−1 or 1 corresponded to the maximum response and 0 corresponded to the baseline response). Neurons whose mean response was less than the baseline were considered as suppressive neurons (n = 13, 9.4%), and the sign of their normalized firing rates was inverted before calculating population-averaged response. Spike trains were smoothed by convolution with a Gaussian kernel (σ = 10 ms) to obtain spike density functions (SDF) for each stimulus. SDF was normalized as the normalized firing rate response, by subtracting the baseline response and then dividing it by the absolute difference between the baseline and maximum response of the neuron. Face selectivity index (FSI) (*43, 44*) was calculated from the mean baseline-subtracted responses to familiar faces (Xfaces) and familiar nonface images (X_nonface_) as: FSI = (X_faces_ -X_nonface_) / (|X_faces_| + |X_nonface_|). In case FSI>1, FSI =1. In case FSI<-1, FSI =-1. FSI = -FSI when both X_faces_ and X_nonface_ are negative, to incorporate inhibitory face-selective response. FSI varies between −1 and 1. When the FSI of a neuron was larger than 0.33, the neuron was considered as face selective.

Time constants of response changes of each neuron were calculated for each stimulus by fitting the responses over the series of experimental session with an exponential function: y = a + b∙e_(c-1)x_, where x is the day after new stimuli introduced, y is firing rate and e is Euler’s number. We used exponential function since the response change is expected to reach a stable value at some time point, and it actually fit well with many of the changing responses (e.g. Fig. 2E). The optimal modeling parameters for a, b and c were estimated by least-squares technique with maximum of 2,000 iterations, within a pre-defined limited range to avoid overfitting: [-500 to 500] for a, [-1000 to 1000] for b and [-10 to 10] for c. Then we defined the time constant τ as |1/c|, which corresponds to the time point when the neuron’s response decreased to 1/e, or 36.8 %, of the response on the first day.

For the population-averaged plot of normalized firing rate, we performed the analysis for neurons which was recorded at least for 70% of total length of evaluated period. For the longer recording sessions of both animals (sessions M-2 and W-2, Supplementary Fig. 2), 54 neurons (81.8%) out of 66 neurons fulfilled this criterion for the first 28 days of recording (Figure 2B-D). We limited our analysis of the time constant τ only for good fitting of exponential function: Exponential fitting was evaluated if R^2^ value of the fitting is larger than 0.3 (42.3% of all fitting), and if the neuron is recorded at least 7 days during first 10 days (95.0% of all fitting), and if the neuron’s response firing rate to the stimulus is larger than 2 Hz (78.4% of all fitting). In addition to the criteria above, we applied two more criteria to evaluated τ only when the neuron’s response changes during the experimental period: we evaluated τ if the neuron’s firing rate is significantly different between first and last two days by t-test (p<0.05, 24.9% of all fitting), and if τ is smaller than 30 days. The second criteria excluded fitting with very long τ which did not show meaningful change of response (254 (3%) cases whose τ was 884.7 ± 637.7 days). With the criteria above, 1377 (16.5%) fitting of 8340 neuron × stimulus combinations (139 neurons × 60 stimuli) were considered (Fig. 2 E, F, Fig.3C, Fig. 4). Of 139 neurons, 124 (89.2%) have at least one fitting to a stimulus and was considered (Fig. 4B). For the time constant τ_trial_, R^2^ value threshold of 0.05 was used instead of 0.3, since the data is not averaged each day and thus noisier than that of τ_day_. For Fig. 3D, if the neurons have at least 10 stimuli which fulfilled the above criteria of exponential regression in both N-10 and N-40 stimulus sets, τ values were averaged across stimuli and used for the analysis of the scatter plot (n=56 for τ_day_ and n=32 for τ_trial_).

## Acknowledgments

We would like to thank David Yu, Charles Zhu and Frank Ye for assistance with fMRI scanning, Katy Smith, George Dold and David Ide for technical assistance and development, Olga Dal Monte and Carlos E. Thomaz for providing face stimli, Vasileva Liubov, Igor Bondar and David Godlove for helpful discussions. Functional and anatomical MRI scanning was carried out in the Neurophysiology Imaging Facility Core (NIMH, NINDS, NEI). This work utilized the computational resources of the NIH HPC Biowulf cluster http://hpc.nih.gov). K.W.K. was supported by the fellowship from the Uehara Memorial Foundation.

## Author contributions

Conceptualization: KWK, DAL Methodology: KWK

Investigation: KWK, EME, JJH, ENW

Data Curation: KWK, EME, JJH, ENW

Formal Analysis: KWK

Visualization: KWK, DAL

Writing – original draft: KWK, DAL

Writing – review & editing: KWK, EME, JJH, ENW, DAL

Funding acquisition: DAL

Project administration: KWK, DAL

Supervision: DAL

---

**Figure S1.**
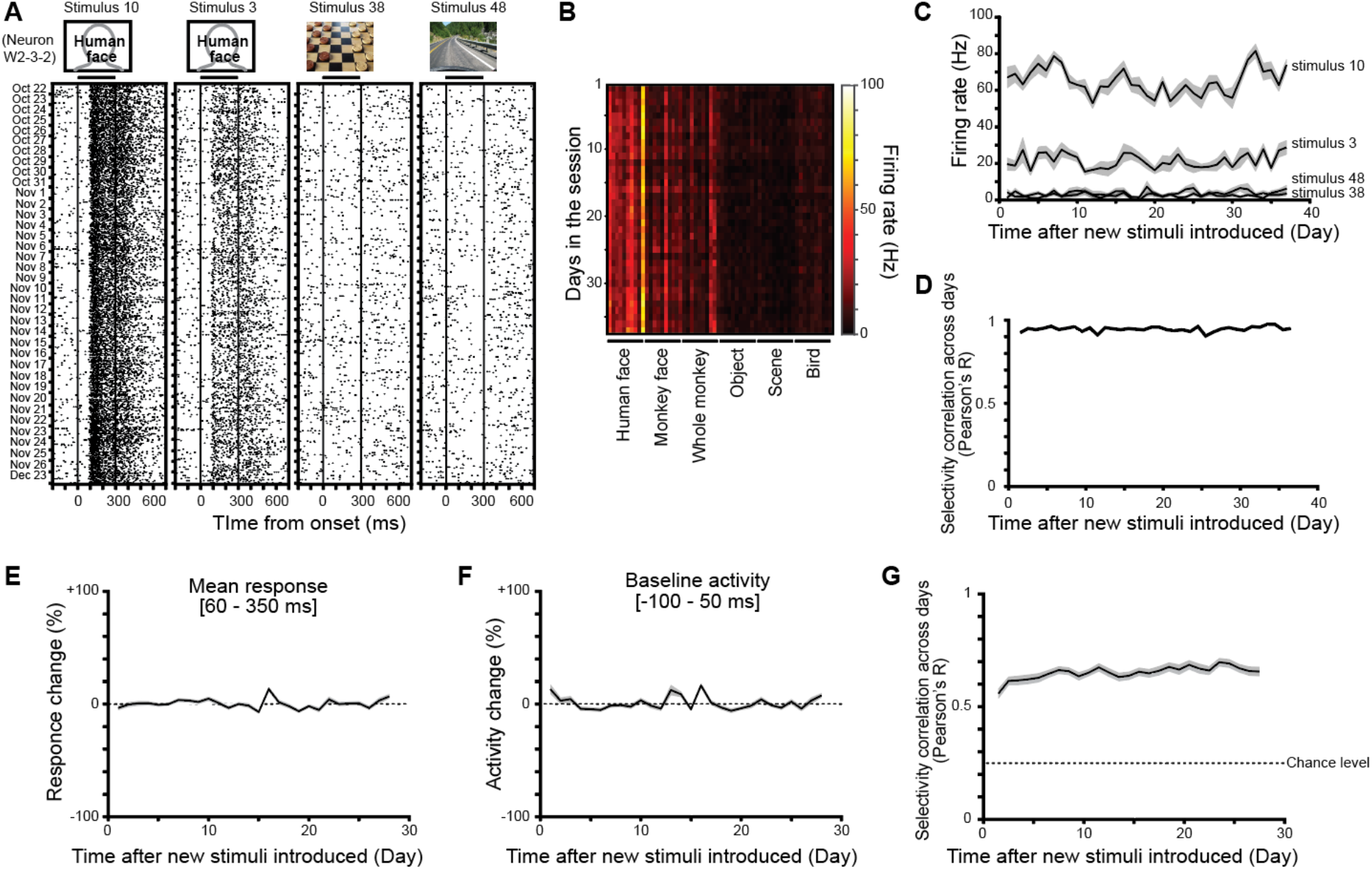
Stable response to familiar stimuli across days. **A**, Responses of an example neuron (neuron W2-3-2) to the familiar stimuli, which is consistent over the recording period of a month. Human face images are substituted to line drawing, according to bioRxiv policy. **B**, Response of the neurons in **A** to all the 60 familiar stimuli, whose response pattern was preserved over the recording period of 37 days. **C**, Firing rate response of the neuron shown in **A** to some of the familiar stimuli over the recording period. **D**, Correlation of the response pattern of the neuron shown in **A**, between two consecutive recording days. **E**, Population mean response to the familiar stimulus set. Ordinate indicates the response deviation from the mean, which is stable across days. There was no significant difference between first and last two days (p=0.34, t_135_=0.95, paired t-test). **F**, Population mean firing rate during baseline period, which is stable across days. There was no significant difference between first and last two days (p=0.07, t_138_=1.84, paired t-test). **G**, Population average of correlation of the response pattern, which is well above the chance level during the recording session There was no significant difference between first and last two days (p=0.27, paired t-test, t_93_=1.11). N=139 for **E**-**F**. N=94 for **G**, which does not include session W-1 where familiar stimuli were presented only on the first and last day of recording.

**Figure S2.**
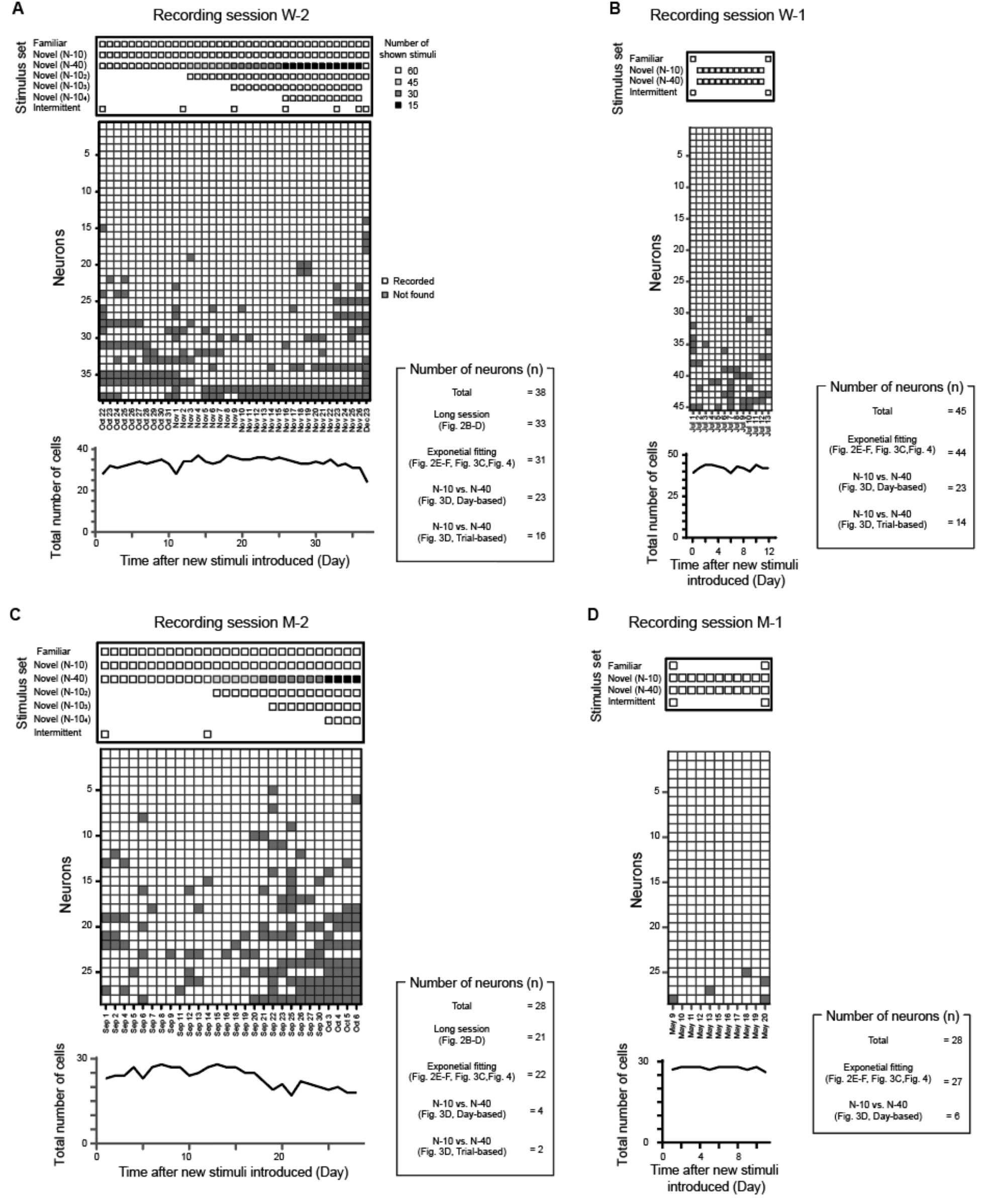
Schedule of stimulus presentation and recorded neurons for each recording session. **A**, recording session W-2. **B**, session W-1. **C**, session M-2. **D**, session M-1. Top, stimulus presentation schedule. Middle, recorded neurons on each day. Bottom, total number of neurons recorded on each day. Box on the right depicted the number of neurons used for each analysis.

**Figure S3.**
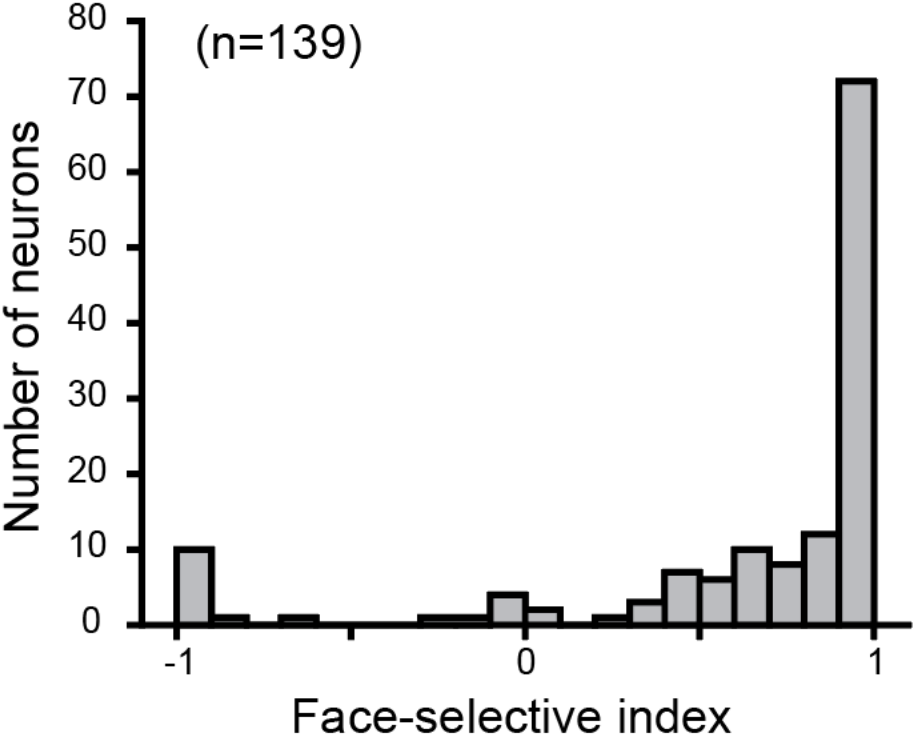
Distribution of face-selective index.

**Figure S4.**
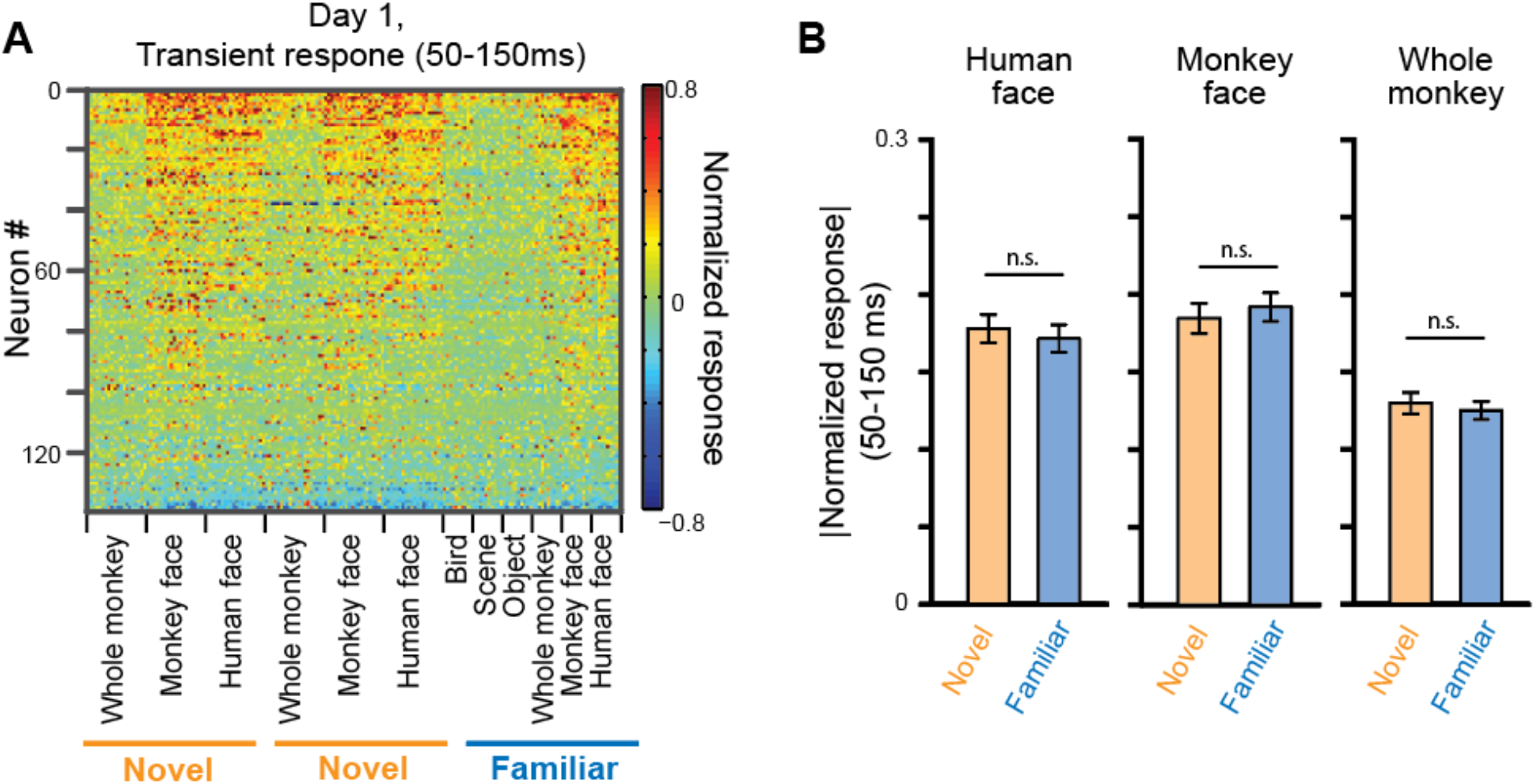
Early transient response on the first day. **A** and **B** are same as **Fig. 1F** and **H**, but for earlier transient response during 50-150 ms.

**Figure S5.**
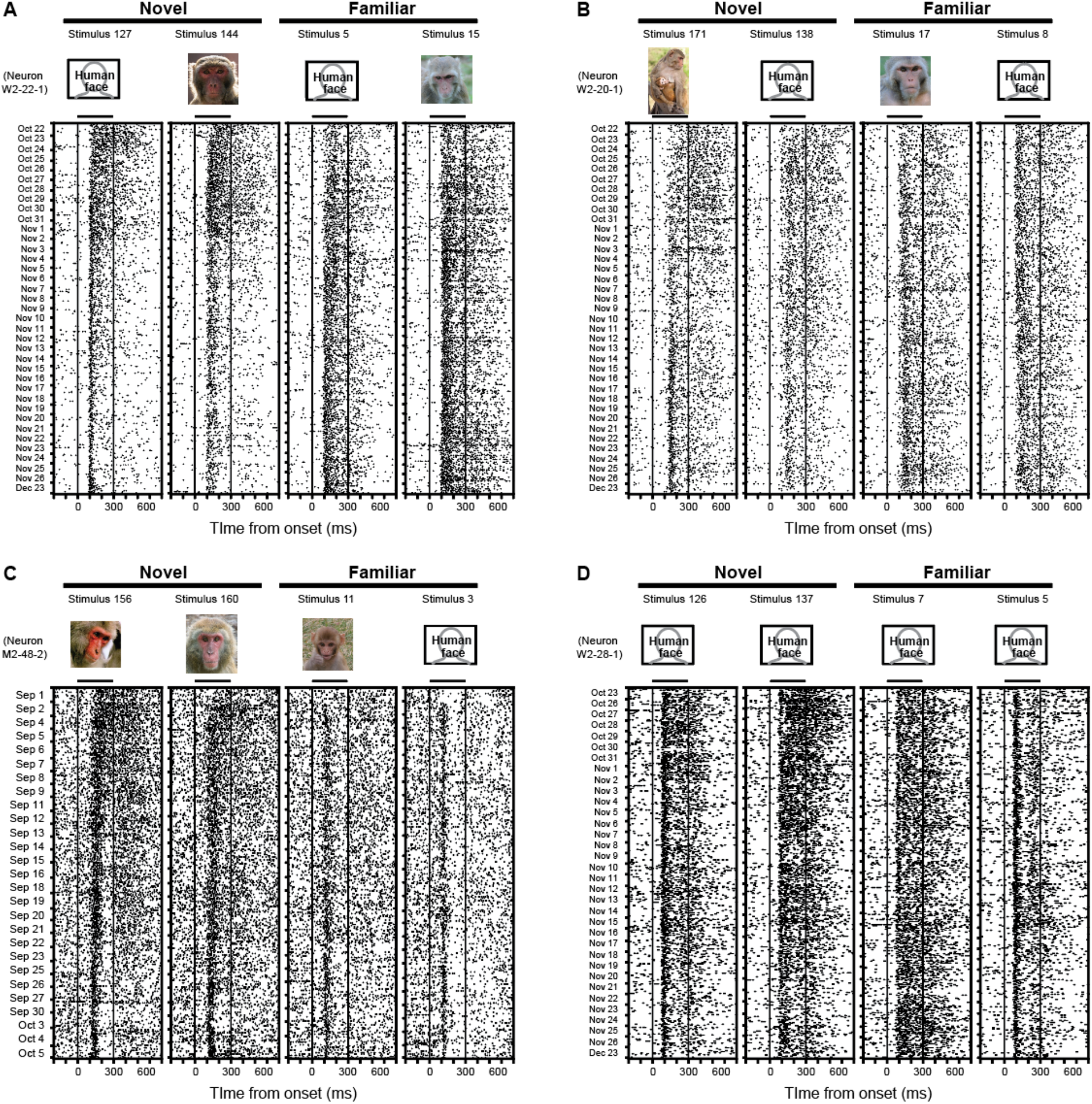
Additional examples of recorded neurons. The neurons showed response decrease in later sustained period for novel stimuli but not for familiar stimuli. Human face images are substituted to line drawing, according to bioRxiv policy.

**Figure S6.**
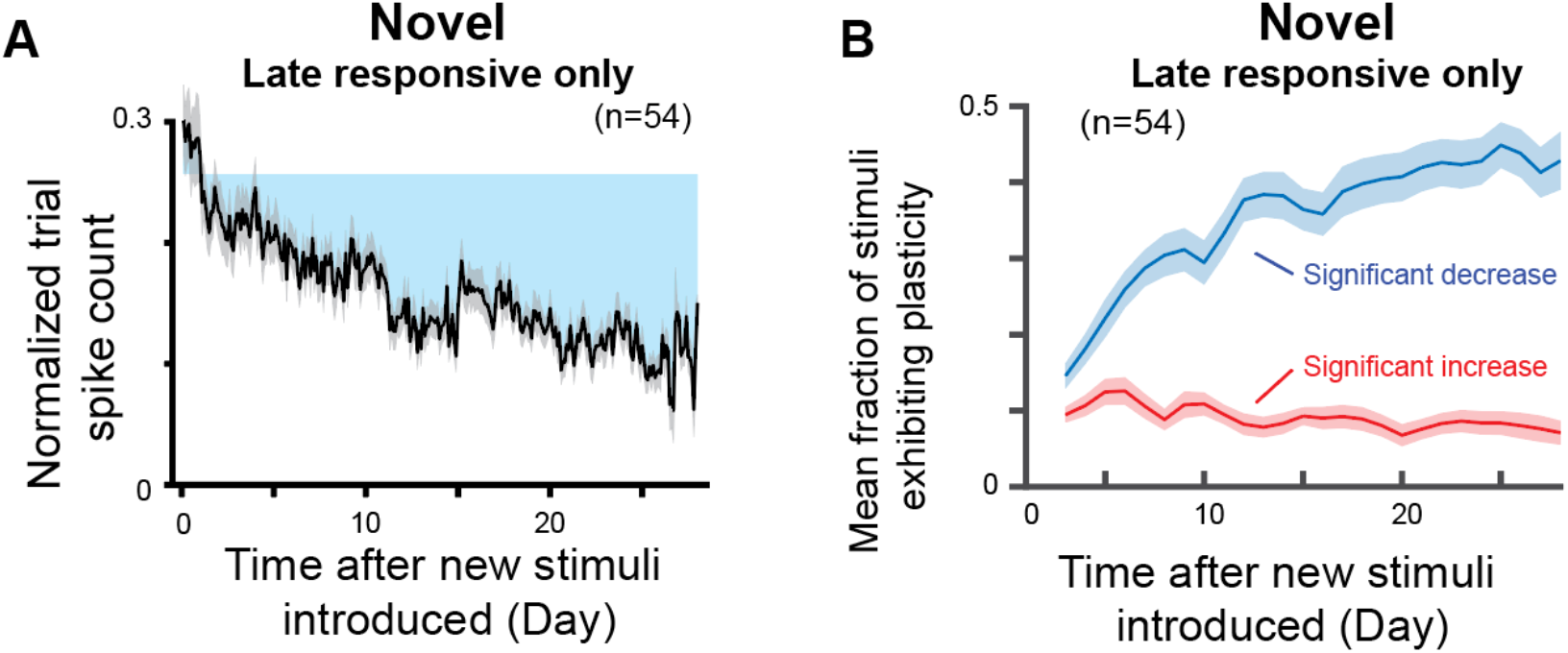
Change of responses to novel stimuli which elicited late sustained response. **A** and **B** are same as **Fig. 2C** and **D** but calculated only for the stimuli which showed significantly higher late sustained response than baseline (p<0.05, t-test with Bonferroni correction).

**Figure S7.**
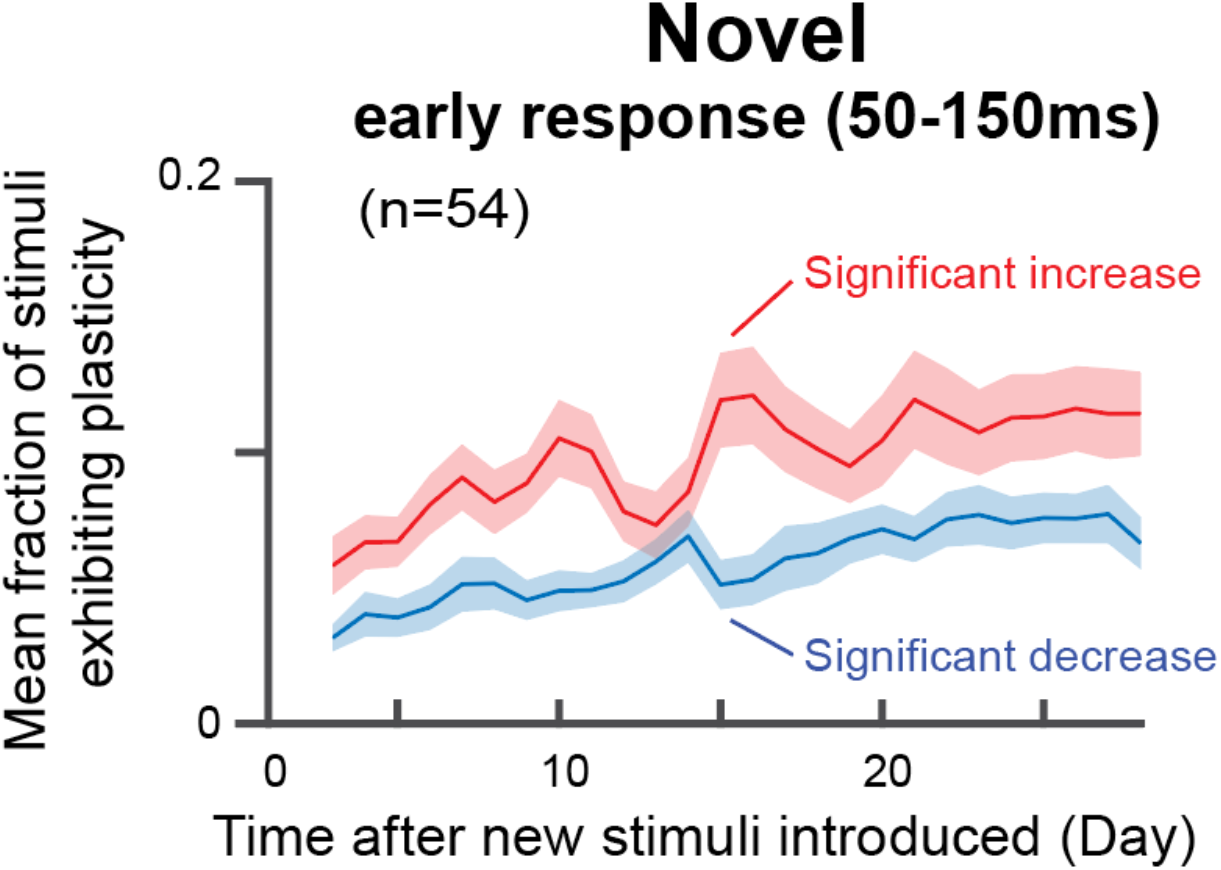
Mean fraction of stimuli which showed significant change of early transient response from that on first two days (t-test, p<0.05). Details are same as **Fig. 2D** but for earlier transient response during 50 to 150 ms after stimulus onset. There was a tendency of slight increase over time in the number of stimuli which showed significant increase.

**Figure S8.**
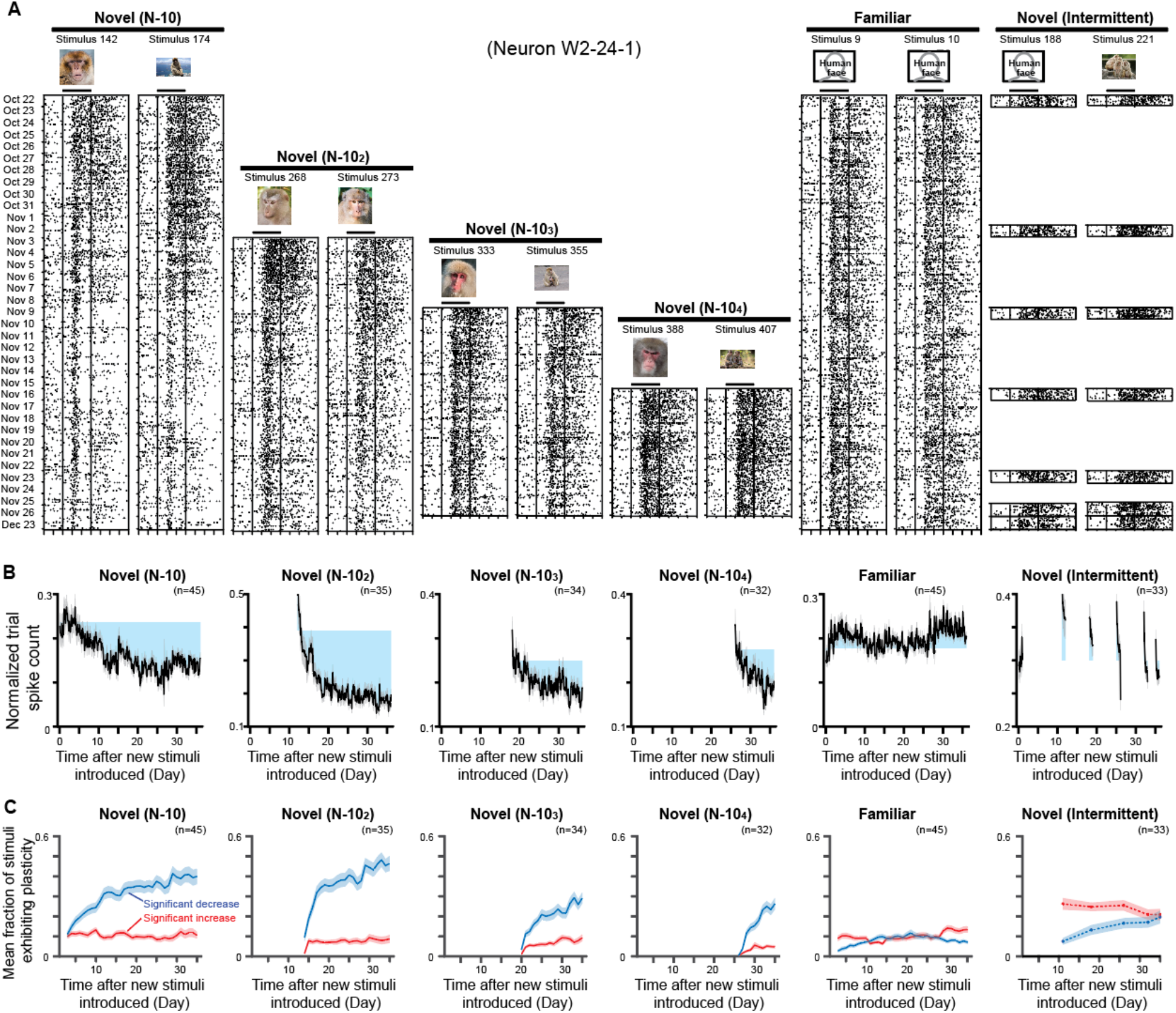
Response decreases for newly introduced novel stimuli at the middle of the recording session. **A**, example response of a neuron (Neuron W2-24-1). Human face images are substituted to line drawing, according to bioRxiv policy. **B**, Change of response during late sustained period over four weeks. **C**, Mean fraction of stimuli which showed significant change from first two days (t-test, p<0.05). **B** and **C** included neurons which was isolated at least 70% of the first 36 days (26 days) of recording. Details of **B** and **C** are same as **Fig. 2C** and **D**.

**Figure S9.**
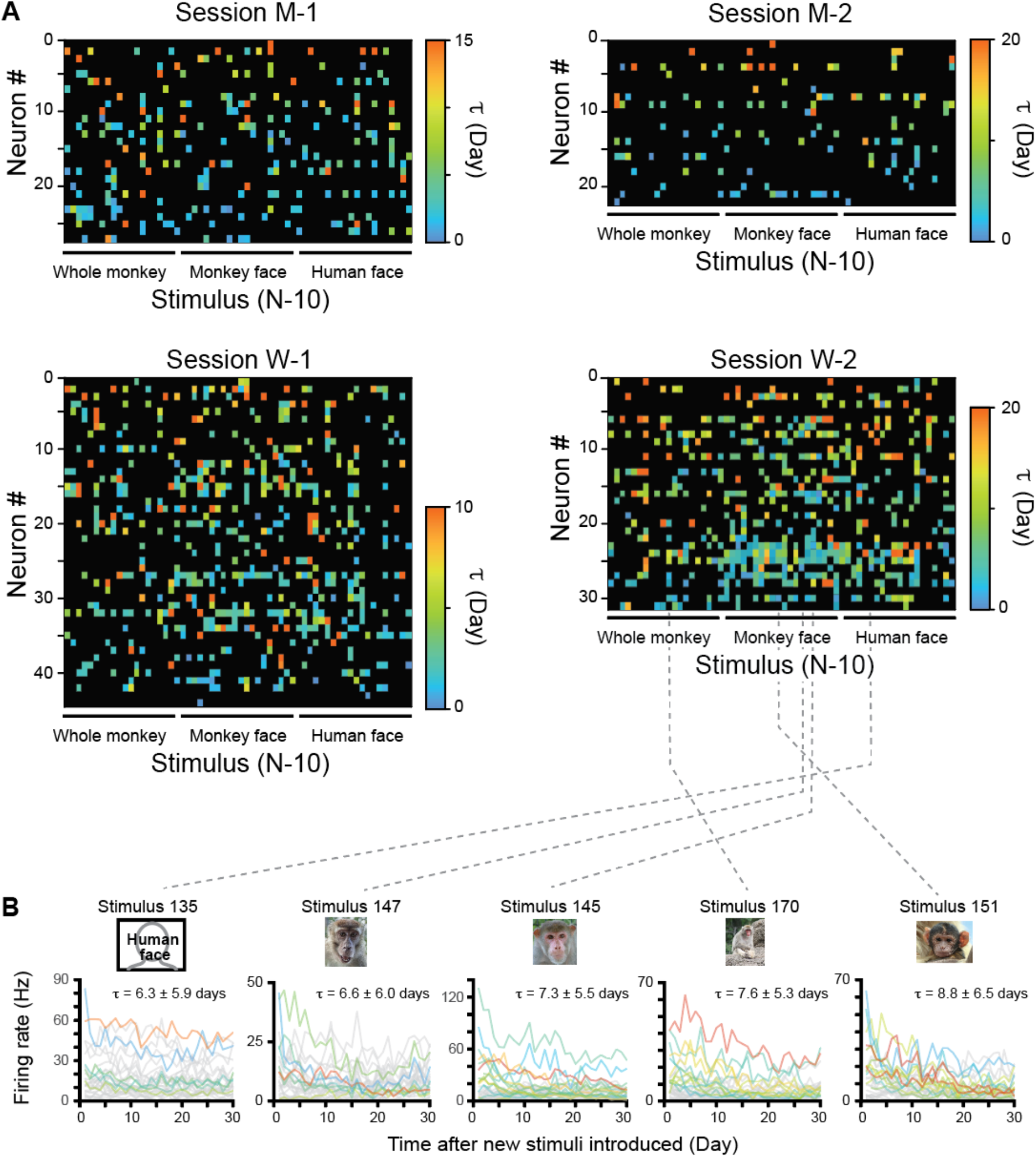
Tau for each recording session. **A**, Time constant for all cell x stimulus combinations of each session. Details are same as **Fig. 4B** but plotted for each recording session. Main effect of cell: session M-1, p<0.01; session M-2, p<0.0001; session W-1, p<0.02; Session W-2, p<10^−8^ (Kruskal-Wallis test). Main effect of stimuli: session M-1, p=0.5; session M-2, p=0.66; session W-1, p<0.03; Session W-2, p<10^−5^ (Kruskal-Wallis test). **B**, Example stimuli of session W-2, showing large variety of plasticity time constants for each stimulus. Details are same as **Fig. 4A** but plotted for each stimulus. Each trace represents response time series of each neuron for the stimulus (n=31 neurons per plot). Human face images are substituted to line drawing, according to bioRxiv policy.

